# Calcium-triggered DNA-mediated membrane fusion in synthetic cells

**DOI:** 10.1101/2023.05.06.539684

**Authors:** Yen-Yu Hsu, J. Chen Samuel, Julio Bernal-Chanchavac, Bineet Sharma, Hossein Moghimianavval, Nicholas Stephanopoulos, Allen P. Liu

## Abstract

In cells, membrane fusion is mediated by SNARE proteins, whose activities are calcium-dependent. While several non-native membrane fusion mechanisms have been demonstrated, few can respond to external stimuli. Here, we develop a calcium-triggered DNA-mediated membrane fusion strategy where fusion is regulated using surface-bound PEG chains that are cleavable by the calcium-activated protease calpain-1.

---

Membrane fusion is the key mechanism that underpins exocytosis and endocytosis, both of which are essential for cell-cell communication in multicellular organisms and are responsible for important biological functions such as synaptic transmission in neurons,^1^ insulin release by pancreatic beta cells,^2^ and granule release by leukocytes and platelets.^3^ In cells, membrane fusion is mediated by coiled-coil interactions between proteins containing SNARE motifs on opposing membranes, which help overcome the kinetic and energetic barriers to induce fusion. A key feature of SNARE-mediated membrane fusion is its calcium dependency, which enables precise spatiotemporal control over exocytosis.^1^

Despite the elegant mechanism of SNARE-mediated fusion, SNARE motifs require multiple post-translational modifications^4^, which introduces challenges for *in vitro* reconstitution. While SNARE-mediated membrane fusion has been demonstrated in a synthetic system,^5,6^ *in vitro* membrane fusions have utilized artificial fusion strategies, including peptide-mediated^7–9^ and DNA oligonucleotide-mediated^10,11^ approaches. Most peptide-mediated membrane fusion designs contain the minimal peptide sequence required to induce membrane fusion from the SNARE motif,^7^ which are commonly denoted as peptide E and peptide K. On the other hand, modifying the vesicles with complementary single stranded DNA can bring opposing membranes into close proximity with each other enabling favourable conditions for fusion.^10^ While robust, both strategies on their own allow fusion to occur spontaneously, with no mechanisms to regulate the process through external stimuli. In addition, few studies have demonstrated spatiotemporal control over membrane fusion by exploiting specific external signals, such as presence of certain ions or pH levels,^9^ to trigger membrane fusion. Crucially, the calcium dependent nature of SNARE-mediated membrane fusion hasyet to be demonstrated in a synthetic system. Incorporating a stimulus-responsive membrane fusion mechanism in synthetic cells would enable the next generation of smart drug delivery vehicles. Consequently, we devise a calcium-triggered membrane fusion strategy, building upon a previously developed DNA-mediated membrane fusion strategy.^10^

Our general methodology is to shield opposing vesicle membranes with surface-bound polyethylene glycol (PEG) chains for blocking membrane interactions in the absence of calcium. These surface-bound PEG chains physically prevent complementary DNA oligos on each surface from hybridizing. A cleavage site for the protease calpain-1 is placed between the membrane-facing cholesterol and the outward-facing PEG chain (Fig. 1). Since calpain is calcium-activated, in the presence of calcium ions, the PEG chains are cleaved off from vesicle membranes by the activated calpain, abrogating their shielding effect and resulting in the exposure and interaction of the complementary DNA oligos for membrane fusion (Fig. 1). To validate this strategy, we started by establishing robust DNA-mediated fusion using cholesterol-conjugated single-stranded DNA oligo pairs developed in Peruzzi *et al*.^10^ Membrane fusion was first tested between supported bilayers with excess membrane reservoir (SUPER) templates and small unilamellar vesicles (SUVs) (Fig. 2A). DNA oligo strands A or B decorate SUPER template membranes and strands A’ or B’ decorate SUV membranes with ∼200 oligos per SUV (ESI†). With strand A on SUPER templates and stand A’ on SUVs, localization of SUV with fluorescently labelled lipids on SUPER templates membranes were observed across 92.8 ± 2% of near all fully formed SUPER templates (Fig. 2A, Fig. S1A, ESI†), suggesting membrane interactions between SUVs and SUPER templates. Similar interactions were observed between SUPER templates and SUVs when strands A and A’ were exchanged for strands B and B’ (Fig. 2B). Conversely, with oligos (strand A) only on the membranes of SUPER templates, no co-localization of SUVs on SUPER template membranes was observed in the vast majority of cases (Fig. 2B), which indicates membrane interactions between SUVs and SUPER templates are DNA-mediated. Furthermore, when SUV membranes were functionalized with strand A, while SUPER template membranes were functionalized with strand B, no interactions between SUVs and SUPER templates were observed in the vast majority of cases (Fig. 2B), which confirms the binding specificity of the complementary oligo pairs.

**Fig. 1.**
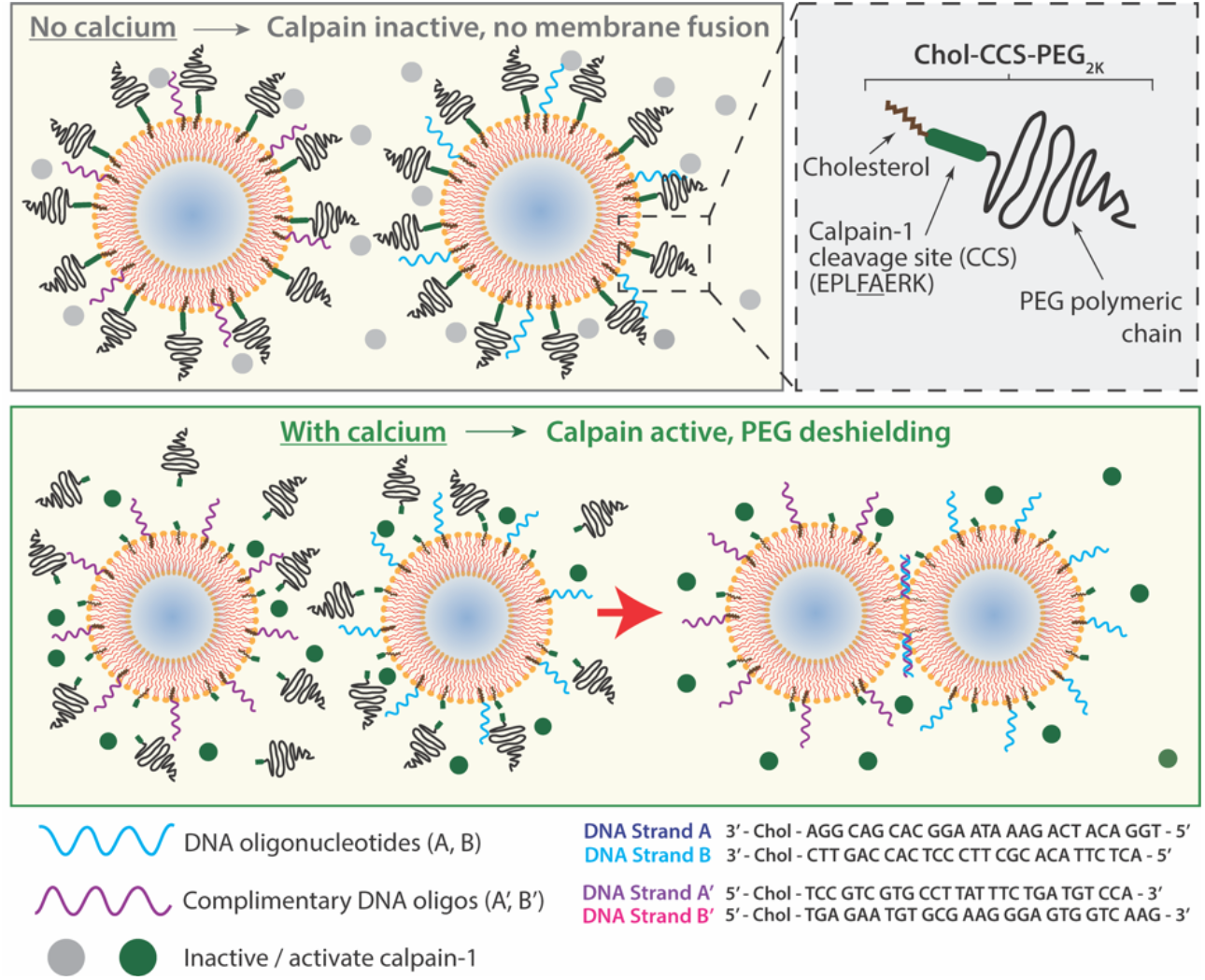
Schematic of DNA-mediated membrane fusion induced by calcium. Complementary single-strand DNA oligonucleotides form double helices in a minimal model for membrane fusion. Without calcium, surface-bound PEG chains linked with a calpain cleavage site physically prevent DNA oligos on opposing membranes from interacting with their complementary DNA oligos. In the presence of calcium, protease calpain-1 activates and cleaves off the PEG chains, which then allows bundle formation between complementary oligos on opposing membranes mediating membrane fusion.

**Fig. 2.**
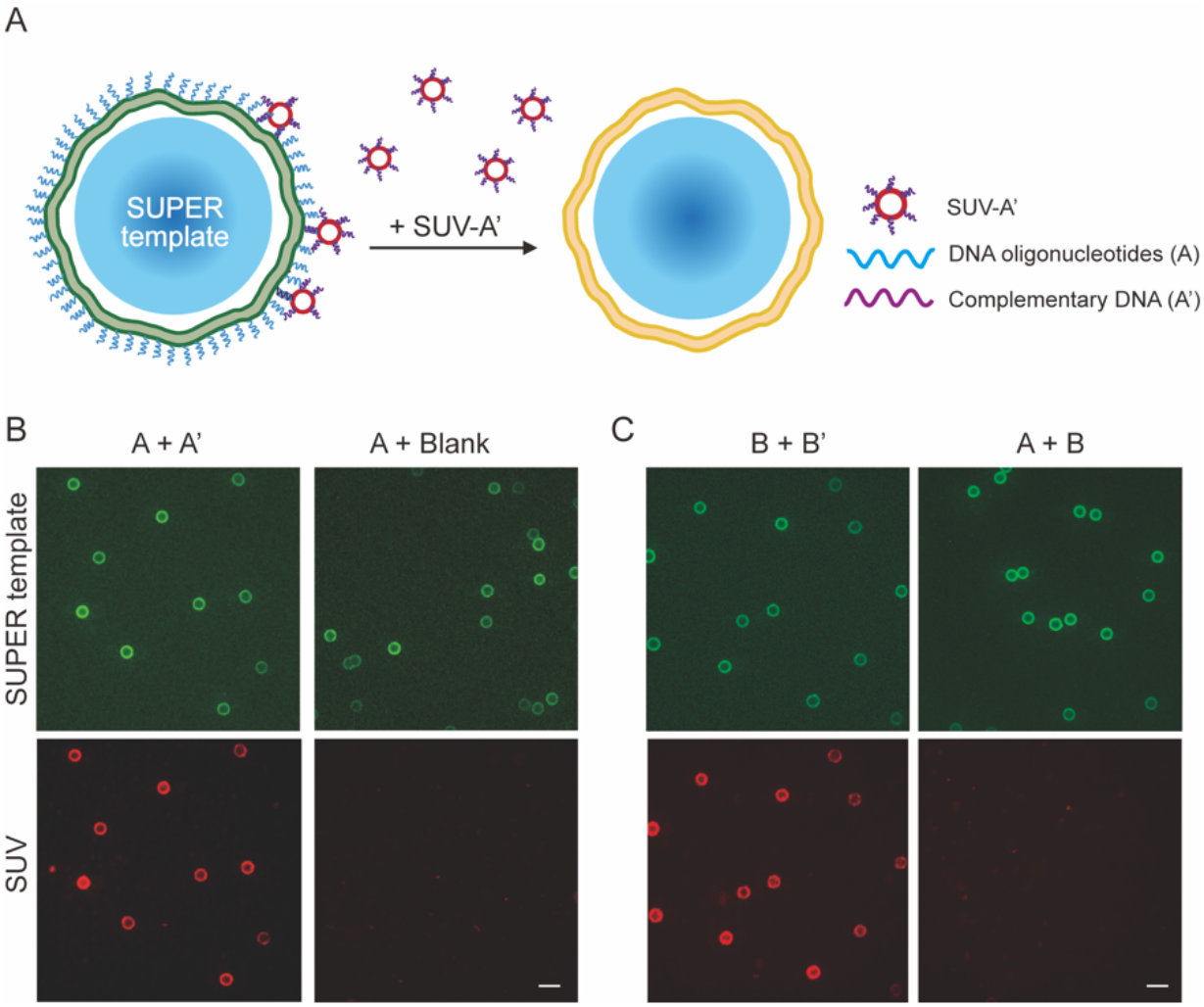
Cholesterol-conjugated DNA oligos mediate membrane interactions between SUVs and SUPER templates. (A) Schematic of DNA-mediated interactions between SUVs and SUPER templates. When mixed together, interactions occur between SUPER templates decorated with strand A and labelled with green fluorescence and SUVs decorated with DNA strand A’ and labelled with red fluorescence. (B) SUPER templates functionalized with strand A interact with SUVs functionalized with strand A’, but not with SUVs without strand A’. (C) Different oligo pair strands B and B’ also induce interactions between SUPER templates (labelled with NBD-PE, green) and SUVs (labelled with Rhod-PE, red) (i.e., strand B on SUPER template and strand on SUV. No membrane interactions between with mismatched oligos (i.e., strand B on SUPER templates and strand A on SUVs). Scale bar: 10 μm.

Following the demonstration of DNA-mediated membrane interactions, we next established the mechanism to regulate these interactions with surface-bound PEG chains. With SUPER templates and SUVs decorated with oligos A and A’, respectively, cholesterol conjugated with PEG chains of various sizes (PEG_1K_, PEG_2K_, and PEG_5K_) were added to SUPER templates and SUVs. We found that SUVs and SUPER templates largely did not interact in the presence of surface-bound PEG_2K_ (Fig. 3A, Fig. S1B, ESI†) or PEG_5k_ (Fig. S1B, S1B, ESI†). However, surface-bound PEG_1K_ was not sufficient to block DNA-mediated membranes interactions (Fig. 3B). Furthermore, having PEG_2K_ on the surface of either SUVs or SUPER templates was also not sufficient to block membrane fusion. In sum, functionalizing opposing membranes with PEG_2k_ is minimally required to prevent membrane interactions mediated by DNA oligos.

**Fig. 3.**
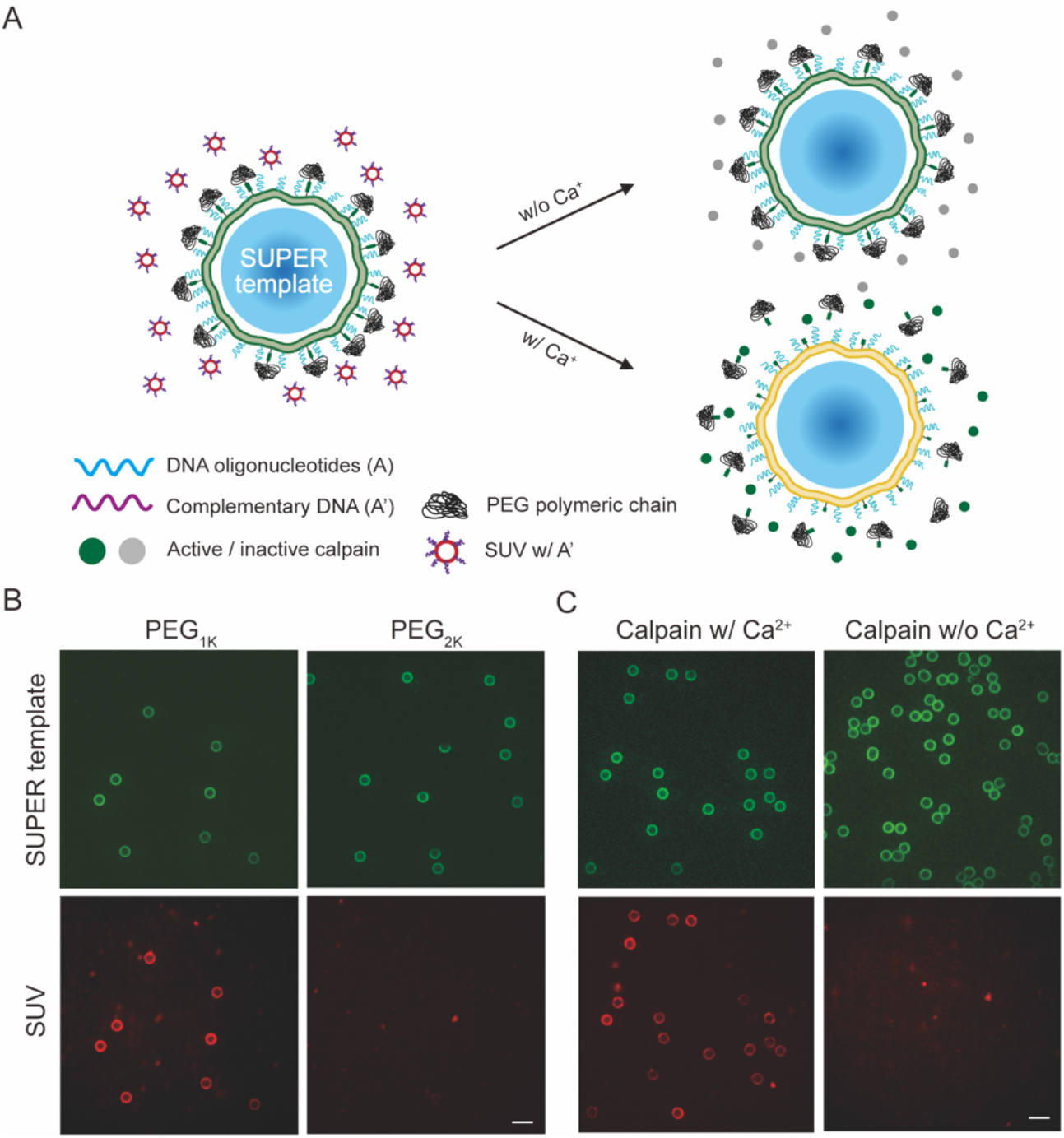
Cholesterol-functionalized PEG chains regulate DNA-mediated membrane interactions. (A) Schematic of calcium dependent regulation of DNA-mediated membrane interactions with Chol-CCS-PEG2K. Similar to Fig. 2, SUPER templates (labelled with NBD-PE, green) and SUVs (labelled with Rhod-PE) are functionalized with complementary DNA oligos of their respective surfaces. (B) Surface-bound PEG2K on both SUPER template and SUV membranes effectively blocks DNA-mediated interactions, but not PEG1K. (C) With Chol-CCS-PEG2K on SUPER template and SUV membranes, membrane interactions are only observed when 225 nM calpain-1 is activated with 5 mM CaCl2. Scale bar: 10 μm.

Next, a custom peptide construct (Chol-CCS-PEG_2K_), denoted as P7 in Table S1, was synthesized where a calpain cleavage site (CCS), EPLFAERK, was conjugated to a cholesterol molecule on one side and PEG_2k_ on the other side. Successful synthesis of the desired peptide (molecular weight: 4.1 kDa) was confirmed by MALDI-TOF mass spectrometry (Fig. S3, ESI†). The ability to cleave CCS was validated first by conjugating it to a fluorescence resonance energy transfer (FRET) dye pair (DABCYL and EDANS).^12–14^ When mixed with calpain-1 and CaCl_2_ in a bulk reaction, the CCS was cleaved, leading to a 2.5-fold increase in EDANS fluorescence over the control condition (Fig. S4A, ESI†). Calpain cleavage was also visualized in an encapsulated system. When calpain FRET reporter, calpain-1, and CaCl_2_ were encapsulated into a giant unilamellar vesicle (GUV), strong fluorescence was observed (Fig. S4B, ESI†). In addition, shuttling calcium across the GUV membrane via ionophores also caused significant EDANS fluorescence in GUVs encapsulating calpain-1 and calpain FRET reporter (Fig. S4B, ESI†). Besides using a calpain FRET reporter, we also confirmed calpain cleavage by gel electrophoresis (Fig. S5, ESI†).

Next, Chol-CCS-PEG2K were added to SUPER template and SUV membranes at a surface density that should induce the PEG chains to exhibit “brush” conformation (ESI†). This will allow PEG to extend further from the membrane surface, potentially enhancing the blockage of fusion of SUPER templates and SUVs concurrently decorated with oligos A and A’, respectively. Minimal membrane interactions were observed when Chol-CCS-PEG_2K_ containing SUPER templates and SUVs were mixed with inactive calpain (Fig. 3C). However, with the addition of 5 mM CaCl_2_ to the mixture, membrane interactions were observed across 85.2 ± 1.8% of fully formed SUPER templates indicating the successful cleavage of membrane-bound PEG chains by calpain-1 (Fig. 3C, Fig. S6, ESI†).

To test DNA-mediated membrane interactions between SUVs and GUVs, we decorated GUV membranes with strand A or B, and SUV membranes with strand A’ or B’. DNA-mediated membrane interactions were observed consistently on the surfaces of 80.9 ± 5.7 % (A+A’) and 77.6 ± 6.8 % (B+B’) of the observed GUVs (Fig. 4A, Fig. S7A, ESI†). Consistent with the earlier result, both PEG_2K_ and PEG_5K_ effectively blocked the DNA-mediated interactions between GUVs and SUVs, while PEG_1K_ did not (Fig. 4B, Fig. S7B, ESI†). To test calpain-mediated membrane fusion, Chol-CCS-PEG_2K_ was incorporated into DNA-decorated GUV and SUV membranes. When GUVs were combined with SUVs and inactive calpain-1, minimal interactions were observed (Fig. 4C). However, with the addition of 5mM CaCl_2_, membrane interactions between GUV and SUVs were observed across 75.8 ± 5.5% of imaged GUVs (Fig. 4C, Fig. S8A, ESI†).

**Fig. 4.**
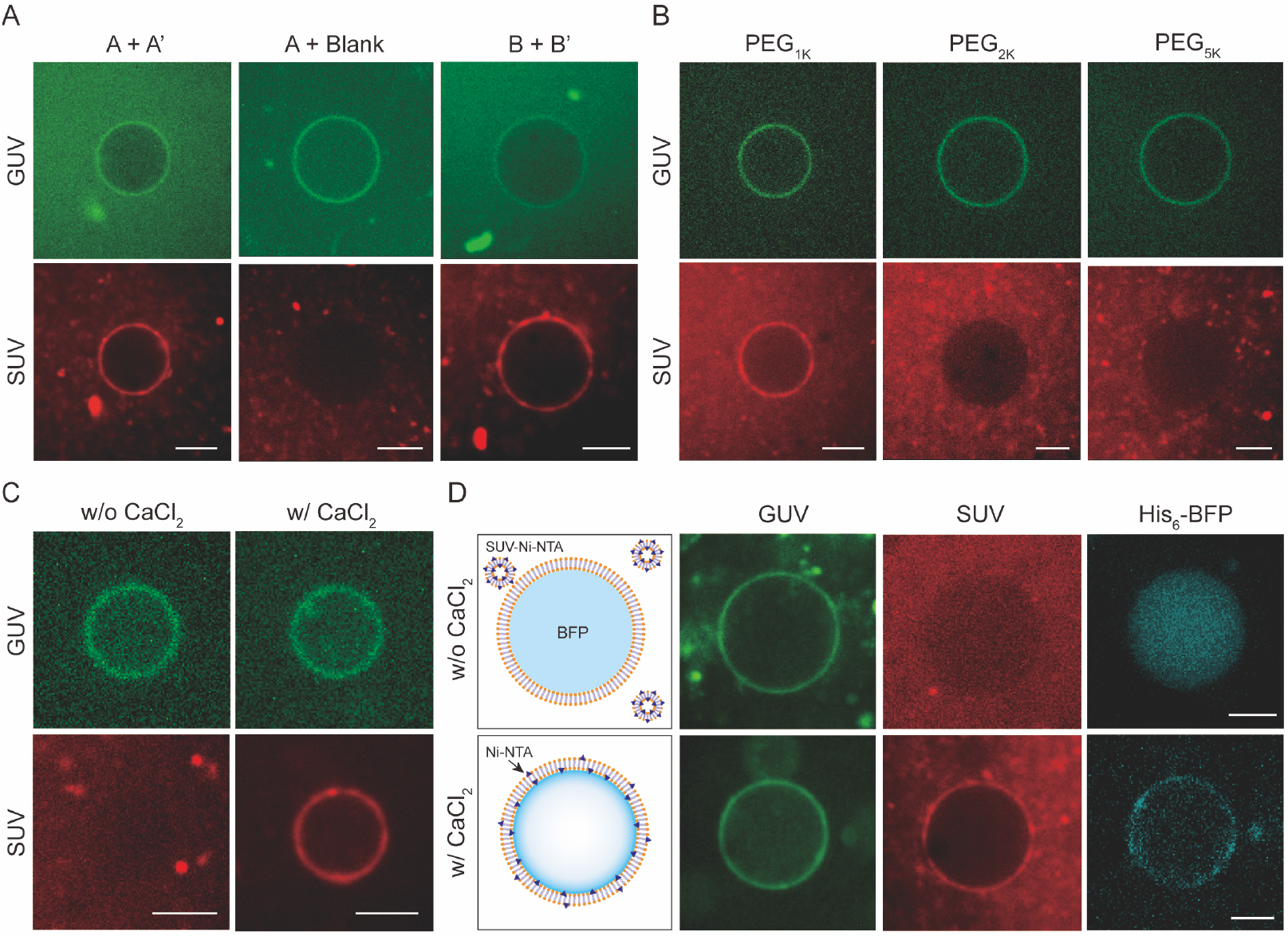
Chol-CCS-PEG2K enables calcium-triggered, DNA-mediated membrane fusion between SUVs and GUVs. (A) Membrane interactions between GUVs (labelled with NBD-PE, green) and SUVs (labelled with Rhod-PE, red) only occur when opposing membranes are decorated with complementary DNA oligos (i.e., GUV with strand A or B, and SUV with strand A’ or B’). (B) PEG2K¬ or PEG5K on both GUV and SUV membranes effectively blocks DNA-mediated interactions, while PEG¬1K does not. (C) Calcium regulates DNA-mediated membrane interaction between GUVs and SUVs, in the presence of calpain-1. (D) Membrane fusion between GUVs and SUVs. SUVs made with Ni-NTA and decorated with Chol-CCS-PEG2K and strand A’ are mixed with GUVs encapsulating His6-BFP and decorated with Chol-CCS-PEG2K and strand A and in the presence of 225 nM calpain-1, with and without 5 mM CaCl2. Scale bars: 10 μm.

However, with the goal of calcium-triggered membrane fusion, the current experiments are not sufficient to prove the fusion of GUV membranes with SUV membranes. Due to the small size of SUVs, which are below the diffraction limit of confocal microscopy, co-localization of SUV membrane fluorescence with SUPER template or GUV membrane fluorescence does not necessarily indicate membrane fusion. Thus, additional experiments like content mixing assays have been used to prove membrane fusion.^7,8,10^ Content mixing assays seek to find evidence that contents encapsulated in SUVs can be delivered and found inside GUVs. However, the detection of content mixing can be challenging, especially since most content-mixing assays rely on fluorescence readouts. This is largely due to the huge volume disparity between GUVs and SUVs; furthermore, the process for generating SUVs leads to the same contents inside and outside the vesicles. Thus, encapsulating SUVs with fluorescent dyes like calcein or fluorescein will require the removal of the dye in the solution containing SUVs, which might be not complete. Thus, the high background fluorescence caused by incomplete dye removal dominates the already weak fluorescent signal inside of GUVs, which results from the substantial dilution of dye when SUVs fuse with GUVs that are approximately a million times larger by volume.

To combat these detection challenges, we developed a novel membrane fusion assay based on the localization of an encapsulated fluorescence protein. Here, SUVs are made with a lipid composition containing DGS-NTA-Ni, a synthetic diacyl lipid carrying a nickel ion with a His-tag binding head group. GUVs encapsulating a histidine-tagged blue fluorescent protein (His_6_-BFP) are prepared by an inverted emulsion-based method,^15,16^ which ensures the contents inside GUVs are completely separate from the outside. When SUVs fuse with GUVs, the Ni-NTA in SUVs should incorporate into GUV membranes, causing the translocation of His_6_-BFP from the GUV lumen to the GUV membrane. When SUVs with strand A’ and Chol-CCS-PEG_2K_ were mixed with GUVs encapsulating His_6_-BFP and decorated with strand A and Chol-CCS-PEG_2K_, BFP fluorescence inside GUVs localized to the membrane, in a manner that is dependent on calpain-1 activity (Fig. 4D). We found 45.7 ± 6 % of GUVs showed membrane interaction with SUVs and BFP translocation to the GUV membrane (Fig. S8B, ESI†). This result strongly supports that DNA-mediated membrane fusion occurs between SUVs and GUVs. The lower efficiency compared with the membrane fusion results in Fig. 3 could be caused by Ni-NTA being positively charged, which limits the amount of Ni-NTA in SUVs. Thus, a significant number of SUVs are required to fuse with each GUV for BFP translocation to be observed. In addition, DNA-mediated membrane fusion is not 100% efficient. Nevertheless, these results demonstrate the initiation of DNA-mediated membrane fusion between SUVs and GUVs in response to calcium-dependent calpain-1 activity.

The strategy for regulating fusion with calpain-cleavable, membrane-bound PEG chains is in theory not limited to DNA-mediated membrane fusion approaches. We believe this strategy can also work for peptide K/E-based membrane fusion.^7,17^ To test this hypothesis, SUPER templates and SUVs were decorated with membrane-bound peptides K and E, respectively. Membrane interactions were only inhibited when surface-bound PEG_2K_ was functionalized on both SUPER template and SUV membranes (Fig. S9, ESI†). While not demonstrated here, we strongly suspect the Chol-CCS-PEG_2K_ construct can also regulate peptide-mediated membrane fusion.

In summary, we have developed a strategy that mimics *in vivo* SNARE-mediated membrane fusion by recapitulating its calcium-dependent nature. Moving forward, this calcium-induced membrane fusion strategy can be integrated into a vesicle-in-vesicle system (i.e., SUVs inside GUV) and functionalized with mechanosensing mechanisms, for example by reconstitution of mechanosensitive channels on GUV membranes.^18^ Such systems would have a significant impact in synthetic cell research by introducing a scheme for controlled synthetic exocytosis only when calcium, a common secondary messenger molecule,^19,20^ is shuttled inside the synthetic cell, triggered by a mechanical stimulus. Such a development would also enable communication between synthetic cells and natural cells in the future.

## Acknowledgement

We thank Dr. Nadab Wubshet (University of Michigan) for providing critical insight and feedback, and Dr. Bineet Sharma (Rutgers University) for discussion. This work is supported by the NIH R01-EB30031 and NSF EF1935265 to A.P.L. and NIH DP2GM132931 to N.S.. A.P.L. conceived the study. Y.Y.H, S.J.C., and A.P.L. designed the experiments. Y.Y.H., S.J.C., J.B.C., B.S., and H.M. performed the experiments. Y.Y.H., S.J.C., J.B.C., H.M., and A.P.L. wrote the paper. All authors contributed to the manuscript revision and approved the final version.

