## Supplemental Information for "Calcium-triggered DNA-mediated membrane fusion in synthetic cells"

### Materials and Methods:

<u>Materials</u>

Fmoc-protected amino acids for peptide synthesis were purchased from EDM Millipore. Fmoc-NH-PEG12-COOH was purchased from Biopharma PEG Scientific Inc. Cholesteryl Hemisuccinate was purchased from Cayman Chemical. Dichloromethane (DCM) was purchased from Millipore Sigma. Dimethylformamide (DMF) and diethyl ether were purchased from Oakwood Chemical Inc. Piperidine was purchased from Alfa Aesar. Oxyma and N,N'-Diisopropylcarbodiimide (DIC) were purchased from ChemImpex. TFA was purchased from Oakwood, and Rink Amide resin was purchased from Novabiochem. All lipids (1,2-dioleoyl-sn-glycero-3-phosphati-dylcholine (DOPC), 1,2-dioleoyl-sn-glycero-3-phosphoethanolamine (18:1 (Δ9-*Cis*) PE (DOPE)), cholesterol, 1,2-dioleoyl-sn-glycero-3-phosphoethanolamine-N-(lissamine rhodamine B sulfonyl) (Rhod-PE), 1,2-dioleoyl-sn-glycero-3-phosphoethanolamine-N-(7-nitro-2-1,3-benzoxadiazol-4-yl) (NBD-PE)) were purchased from Avanti Polar Lipids. Cholesterol-conjugated oligonucleotides were purchased from Integrated DNA Technologies and cholesterol-conjugated PEG chains were purchased from Creative PEGWorks. EDANS/DABCYL FRET reporter for calpain-1 was purchased from GenScript. 16.5% Mini-PROTEAN® Tris-Tricine Gel and Precision Plus TM Protein Unstained Protein Standards were purchased from Bio-Rad Laboratories. All chemicals were purchased from

Sigma unless otherwise noted. Peptides were custom synthesized and listed in **Table S1** (more details below).

Cloning and preparing DNA construct for $His_6$-BFP

A plasmid for bacterial expression of mTagBFP with a C-terminal 6xHis tag was generated by cloning the sequence encoding mTagBFP into a pET28b vector. The backbone fragment was amplified from a pET28b vector (a gift from Tobias Pirzer, Technical University of Munich, Germany) with forward primer: CACCACCACCACCACCAC and reverse primer: AAAAAACCTCCTTACTTTCTAGTCTCAAG. Similarly, the insert fragment was amplified from sTag-BFP plasmid (Addgene #186905)[1] using forward primer: TCTTGAGACTAGAAAGTAAGGAGGTTTTTTATGTACACCATCGTGGAGCAGTAC and reverse primer: AGCCGGATCTCAGTGGTGGTGGTGGTGGTGCAGATCCTCTTCTGAGATGAG. The fragments were assembled using Gibson Assembly, and the cloning sequences were verified using Sanger sequencing (Eurofins).

Bacterial expression and purification

Protein expression and purification from bacteria were performed following the conventional His-purification reported elsewhere[1]. pET28b-mTagBFP construct was transformed into BL21-DE3-pLysS cells (Agilent). A single colony was picked from the LB plate and grown in 5 mL LB supplied with 50 µg/mL kanamycin at 37 ºC shaking at 220 rpm overnight. Next day, the culture was diluted in 1 L LB supplemented with 0.8% w/v glucose and 50 µg/mL kanamycin. The culture was grown at 37 ºC shaking at 220 rpm and was induced with 0.42 mM isopropyl β-D-1-thiogalactopyranoside (IPTG) once the $A_{600}$ reached 0.5-0.6. After induction, the culture was incubated at 30 ºC shaking at 200 rpm for 4-5 h. The cells were then harvested through centrifugation at 5000 g for 10 min. The cell pellet was resuspended in 30 mL of lysis buffer (50 mM Tris-HCl (pH 7.4), 300 mM NaCl, and 1 mM 4-(2-aminoethyl) benzenesulfonyl fluoride hydrochloride (AEBSF)). The cells were then lysed using a tip sonicator (Branson Sonifier 450) and the lysate was centrifuged at 30000 g for 25 min. Next, the supernatant was run through an equilibrated HisTrap column (Cytiva) in an AKTA start fast protein liquid chromatography system. The column was washed with 15 column volumes of washing buffer (50 mM Tris-HCl (pH 7.4), 300 mM NaCl, and 50 mM imidazole) before the purified protein was eluted with 10 column volumes of elution buffer (50 mM Tris-HCl (pH 7.4), 300 mM NaCl, and 400 mM imidazole) and collected in 1 mL fractions. The purification quality was assessed for each fraction by SDS-PAGE, and the fractions with high concentration of the protein were pooled and dialyzed against 1 L PBS overnight at 4 ºC. The protein concentration was measured with NanoDrop (Thermo Fisher Scientific) (extinction coefficients predicted by Benchling) before it was aliquoted and stored at −80 ºC until use.

**Supplementary Information**

Preparation of GUVs

GUVs were generated via electroformation following a published protocol[2] for all membrane interaction experiments, due to the method's ability to reliably produce high GUV yield. GUVs were formed by the droplet transfer approach (i.e., inverted emulsion-based method)[3,4] for all membrane fusion experiments. 0.5 mM lipid dissolved in chloroform was mixed in molar ratios specified in **Table S2** for different experiments. The lipid mixture was evaporated under a stream of nitrogen to remove any remaining chloroform in a glass vial. 1 ml of mineral oil (Sigma-Aldrich) was then added to the glass vial and then mixed with the dried lipid film by vigorous vertexing. The lipid-in-oil solution was placed in the oven at 60°C for 20 minutes. During incubation, the outer solution was prepared by mixing the desired amount of glucose solution into 1X PBS solution to match the osmolarity of the inner solution to be encapsulated inside the GUVs. For membrane interaction experiments, the inner solution was 300 mM sucrose solution in milli-Q water. For membrane fusion experiments, the inner solution contains $His_6$-BFP with 5% OptiPrep. In a 1.5 ml epitube, 300 µl of lipid-in-oil solution was carefully added on top of 400 µl of outer solution and incubated at room temperature for 1 hr to form a lipid monolayer at the interface, followed by carefully adding to the top water-in-oil single emulsion droplets generated by vigorously pipetting 20 µl of inner solution with 600 µl of lipid-in-oil solution. After centrifugation at a speed of 2.5 k RCF for 10 mins, the oil on top was carefully removed and the pellet fraction was gently resuspended and transferred to a new epitube.

Preparation of SUVs

SUVs were prepared by using a thin film hydration method followed by extrusion. 1 mM lipids dissolved in chloroform were mixed in molar ratios specified in **Table S3**. The lipid was dried under vacuum for 1 h to create a uniform lipid film and remove any remaining chloroform in a glass vial. 1 ml of 1X PBS was then added to the film and thoroughly vortexed. The mixture was then passed through a liposome extruder (Avanti Polar Lipids) with a 100-nm porous membrane 15 times to generate SUVs. For calcium-dependent membrane fusion experiment, calpain-1 cleavable PEG chains (Chol-CCS-$PEG_{2K}$) were added to the lipid mixture at a final concentration of 1 mM so the lipid composition becomes DOPC:DOPE:cholesterol:Chol-CCS-$PEG_{2K}$:Rhod-PE/NBD-PE = 63.9:5:29:5:0.1.

SUPER template generation

SUPER templated beads were generated following a published protocol.[5] For SUPER template formation, 25 µl of SUV solution was fused with 5 µl of 5-µm silica beads (Bangs Laboratories) in the presence of 1 M NaCl. The final SUPER templated beads were washed with PBS twice by centrifuging at 200 × g for 2 min and then resuspended in 30 µl of milli-Q water at a final concentration of ~2.4 × $10^7$ beads/ml. The SUPER template stock can be stored at room temperature for 3 hr.

Peptide synthesis and characterization

Peptides mentioned in **Table S1** were obtained using solid phase peptide synthesis (SPPS) on a CEM Liberty Blue instrument. Synthesis was performed on a solid phase Rink-Amide resin (0.78 mmol/g) at a 0.1 mmol scale, using a standard Fmoc protocol and deprotected in 20% piperidine in DMF. Amino acids, PEG blocks, cholesterol hemisuccinate, coupling agents, DIC and Oxyma, were added in a 10-fold molar excess. Crude peptides were cleaved by shaking the resin in a solution containing trifluoroacetic acid (TFA), triisopropylsilane (TIPS), and water in a ratio of 95:2.5:2.5 for 3.5 h. The resin was washed with TFA and concentrated under nitrogen. The solution was then added to 40 mL of cold diethyl ether to precipitate the peptide. The solution was centrifuged at 4200 rpm for 10 min, the supernatant was removed, and the pellet was allowed to dry overnight. The dried pellet was dissolved in a mixture of water and acetonitrile (50:50) and 0.1% TFA. Peptides were purified via reverse phase chromatography on a Waters HPLC using a Phenomenex column with C18 resin. A linear gradient was generated using water/acetonitrile + 0.1% TFA, from 10% to 100% acetonitrile over 50 minutes. Peak fractions were collected based upon their absorbance at 230 nm and tested for purity by MALDI-TOF mass spectrometry on Bruker microflex LRF MALDI using α-cyano-4-hydroxycinnamic acid matrix (Sigma). Pure fractions were pooled and lyophilized, and peptides were stored at -20 °C until use.

Membrane interaction assay

To functionalize vesicles with DNA oligonucleotides, SUVs consisted of 1 mM lipids with mole percentage of 69.9% DOPC, 10% DOPE, 20% cholesterol and either 0.1% Rhod-PE or NBD-PE were functionalized with ~200 oligos per vesicle (calculated following formulas given in Peruzzi *et al.*) [6] by incubating SUVs with 10 μM cholesterol-conjugated DNA strand A or B for 30 mins. SUVs containing NBD-PE were later used for generating lipid-coated beads. Similarly, GUVs were functionalized with 10 μM of cholesterol-conjugated DNA strand A' or B' for 30 mins. To prevent close contact between vesicle membranes, 40 μM of cholesterol-conjugated PEG chains were added to SUVs, GUVs, and SUPER templates and incubated for 30 minutes. For calcium-dependent membrane interaction experiment, a lipid composition of 64.9% DOPC, 10% DOPE, 20% cholesterol, 0.1% Rhod-PE or NBD-PE and 5% Chol-CCS-$PEG_{2K}$ was used instead. For the SUV-SUPER template experiment, SUVs and SUPER templates labeled with complementary DNA oligos were mixed in a 1:4 volume ratio and incubated together for 30 minutes. Afterwards, SUPER templates were washed with 1X PBS twice by centrifuging at 200 g for 2 mins and then resuspended in 30 μl of PBS. The solution containing SUPER template was then deposited onto a coverslip chamber and imaged using a 60x oil objective.

**Supplementary Information**

For the SUV-GUV experiment, SUVs and GUVs labeled with complementary DNA strands were mixed in a 1:4 molar ratio (~0.83 mM final lipid concentration) and incubated together for 30 minutes. The mixture was then transferred to a 96-well glass-bottom plate and imaged on a confocal microscope using a 60x objective.

Membrane fusion assay

To visualize membrane fusion triggered by calcium, GUVs with 94.9% POPC, 5% DOPE, and 0.1% NBD-PE and encapsulating 6.25 μM $His_6$-BFP were made via the inverted emulsion method outlined above. SUVs consisting of 1 mM lipids with 34.9% DOPC, 15% cholesterol, 0.1% Rhod-PE and 50% DGS-NTA (Ni) were extruded 15x through a 400 nm filter. GUVs were functionalized with 10 μM of DNA-oligos while SUVs were functionalized with ~200 oligos per vesicle using the method described above. GUVs and SUVs were mixed in a 1 to 4 molar ratio and incubated for 30 mins. The vesicle solution was then deposited into a 96-well plate chamber and imaged on a confocal microscope using a 60x objective.

Confocal fluorescence microscopy

All images were acquired using an oil immersion 60×/1.4 NA Plan-Apochromat objective with an Olympus IX-81 inverted fluorescence microscope (Olympus) controlled by MetaMorph software (Molecular Devices) equipped with a CSU-X1 spinning disk confocal head (Yokogawa), a custom laser launch with solid-state lasers (Solamere Technology Group), and an iXON3 EMCCD camera (Andor). Images of BFP and lipid fluorescence were acquired with 405-nm and 488-nm laser excitation at an exposure time of 500 ms, and with 561-nm laser excitation at an exposure time of 200 ms, respectively. Each acquired image contained ∼10 lipid bilayer vesicles or ∼10 lipid-coated beads that had settled upon a 96-well glass-bottom plate or a coverslip, respectively. Three images were taken at different locations across a well or coverslip for an individual experiment. Three independent repeats were carried out for each experimental condition. Samples were always freshly prepared before each experiment.

FRET calpain biosensor plate reader assay

250 nM FRET-pair calpain-1 biosensor & 450 nM calpain-1 were combined with 2 μL OptiPrep and 2 μL reaction buffer, which consists of 50 mM KOH & 10 mM dithiothreitol (DTT), and then DI water was added to a total reaction volume of 20 μL. Each reaction is loaded into a 384-well glass-bottomed well plate and the corresponding EDANS fluorescence intensity was measured at 336/455 nm wavelength using a Synergy H1 plate reader (BioTek). After baseline fluorescence measurements, 5 mM $CaCl_2$ was added to the reaction. The reaction is then immediately imaged with the same settings for 1 h.

Bulk reaction & gel electrophoresis of peptides with calpain cleavage site

**Supplementary Information**

Peptides P3 & P6 were taken from the stock solution, which was dissolved in chloroform. Peptides were dried under a stream of Argon gas, and then desiccated for 30 min. Peptides were then rehydrated in reaction buffer to a working concentration of 10 µM. For bulk reactions, a combination of peptides (10 µM), calpain-1 (225 nM), or $CaCl_2$ (3 mM) was mixed to a total reaction volume of 15 µL in a microcentrifuge tube. Reactions were then incubated at 37 ºC for 30 min. After completion, reactions were mixed with tricine sample buffer (1:1 mix ratio) and 2% β-mercaptoethanol, and then heated at 90 ºC for 5 min. SDS-PAGE gels were run in a 16.5% polyacrylamide tris-tricine precast gel at 100V. After electrophoresis, the gel was stained with Coomassie G-250 stain for 1 h on a rocking shaker, and then washed overnight in DI water. Gel images were acquired in a Sapphire Biomolecular Imager (Azure Biosystems).

Image and data analysis

All images were analyzed in MATLAB and in a nonblinded manner. Since all the GUVs were labeled with green fluorescence from NBD-PE, the edges/boundaries of vesicles were first detected and isolated using the built-in MATLAB function “imfindcircles”. The areas that GUVs cover were marked. Next, the red fluorescence images, which portray SUVs, were analyzed. The average background intensity of red fluorescence was measured by averaging the fluorescent intensity measurements of all locations across one image except areas where the GUVs were located. Then, the average red fluorescence along the GUV perimeter (i.e., all pixels that fall on a one-pixel wide line, which traces the GUV perimeter) was compared to the average background intensity. If the red fluorescence along the perimeter was at least 1.2x greater than the background, then co-localization of SUV and GUV membrane fluorescence was observed. Consequently, GUV-SUV interactions were implied. Lastly, the ratio of GUVs that had interactions with SUVs could be quantified by comparing the number of GUVs with GUV-SUV interactions with the total number of GUVs detected.

Taking the GUVs that were identified earlier that had membrane interactions with SUVs, we then quantified if membrane fusion occurred. For each image, we averaged its background fluorescence (blue) using the same method conducted on red fluorescence images. Next, the blue fluorescence on and inside the GUV was subtracted by the average background fluorescence. All the subtracted values were averaged and defined as the average GUV lumen fluorescence. Furthermore, still using the subtracted fluorescent values, the GUV membrane fluorescence was quantified using the method for membrane fluorescence quantification of red (i.e., from SUV) fluorescence. Lastly, the GUV lumen and membrane fluorescence were compared; recruitment of $His_6$-BFP from GUV lumen to membrane was quantified when the GUV membrane fluorescence was at least 2x greater than the GUV lumen fluorescence. By comparing the GUVs with membrane fusion and the ones with membrane interactions, the percentage of GUVs experiencing membrane interactions, as well as membrane fusion could be calculated.

**Supplementary Information**

### Supplemental Calculation

PEG conformation calculation

$\boldsymbol{D}$ is the distance between PEG graft and $\boldsymbol{R_f}$ is the Flory radius of the PEG graft. $\boldsymbol{A}$ is the area occupied per PEG chain, $\boldsymbol{a}$ is the monomer length of the PEG chain (0.35 nm), $N$ is the degree of polymerization (i.e., number of PEG repeats, which is 48 repeats for $PEG_{2K}$). For Chol-CCS-$PEG_{2K}$ construct, assuming the PEG conformation only depends on the PEG chain in the construct, we can use the equations below to compute $\boldsymbol{D}$ and $\boldsymbol{R_f}$ values,[6] which will help us predict the expected PEG conformation on SUV membranes.

$$\boldsymbol{D = 2\left(\frac{A}{\pi}\right)^{\frac{1}{2}}}; \quad \boldsymbol{R_f = aN^{3/5}}$$

Specifically, if the $\boldsymbol{R_f/D}$ ratio is below 1, PEG chains are in the "mushroom" conformation. If $\boldsymbol{R_f/D}$ is greater than 1, PEG chains are in the "brush" conformation.[6] For $PEG_{2K}$, the Flory radius is 3.57 nm. With the addition of 5% PEG by count (mole), and since all the PEG is on the outer membrane PEG will account for 10% of the composition of the outer membrane assuming both inner and outer membrane of the SUV have the same surface area. Based on the 100 nm size of the SUVs, the surface area of SUVs is approx. 31,400 $nm^2$. Assuming that a phospholipid molecule takes up around 65 $Å^2$ (0.65 $nm^2$),[7] each SUV will contain around 48,000 molecules on its outer membrane. Of which, Chol-CCS-$PEG_{2K}$ will account for around 4,800 of them. Thus, on average, each PEG chain will occupy an area of 6.54 $nm^2$. Thus, the distance between PEG grafts can be calculated to be 2.89 nm. As a result, the $\boldsymbol{R_f/D}$ ratio is 1.24, which predicts that PEG exhibits a "brush" conformation.

**Supplementary Information**

**Supplementary Information**

### Supplemental Figures

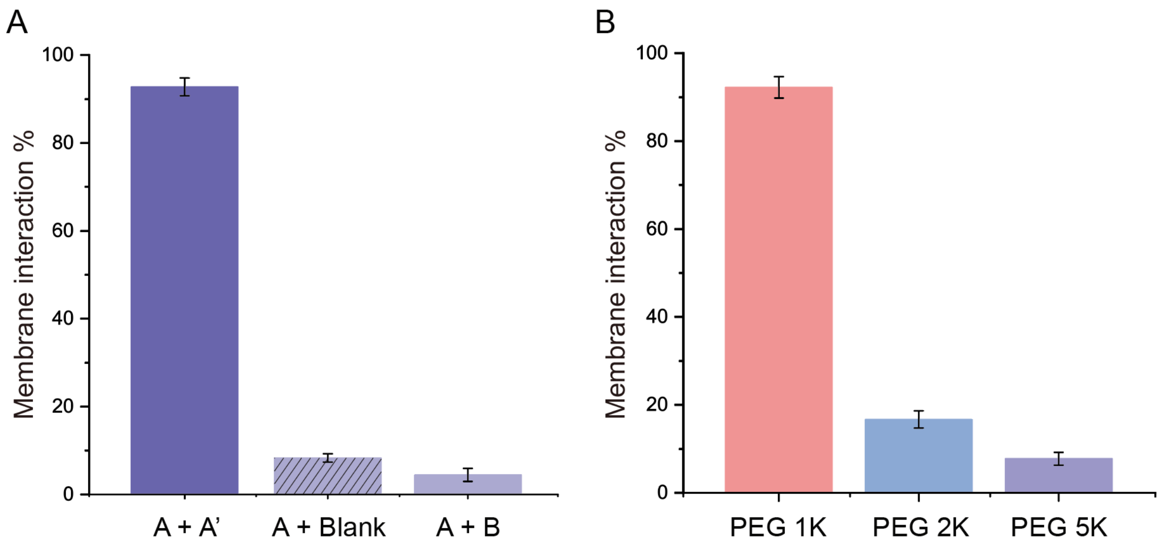


**Figure S1. Quantification of DNA-mediated membrane interactions between SUPER templates and SUVs.** (A) SUPER templates labeled with NBD-PE are functionalized with DNA strand A while SUVs labeled with Rhod-PE are functionalized with either DNA strand A', no DNA, or DNA strand B. The bar graphs represent the percentage of SUPER templates showing membrane interaction with SUVs by the red fluorescence detected on SUPER template membranes. (B) Both SUPER templates and SUVs membranes were covered with complementary DNA and PEG chains. The bar graphs display the percentage of lipid-coated beads showing red fluorescence observed on SUPER templates to the total number of lipid-coated beads. At least 60 lipid-coated beads were analyzed for each condition. All experiments were repeated three times under identical conditions. The error bars represent standard errors.

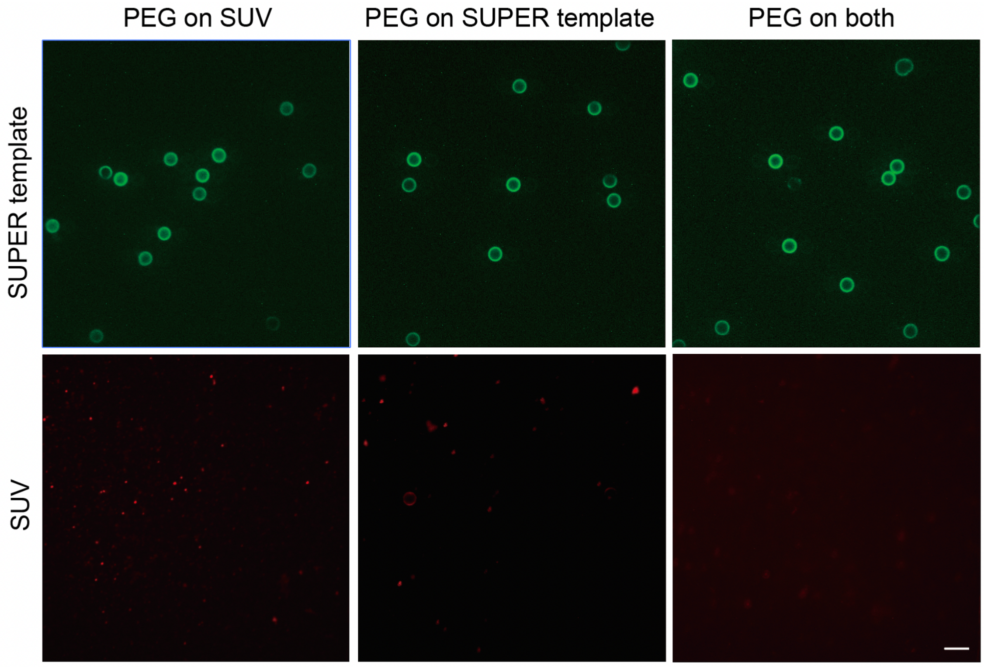


**Figure S2. Blocking DNA-mediated interaction between SUVs and SUPER templates using surface-bound $PEG_{5K}$.** Functionalizing cholesterol-conjugated $PEG_{5K}$ on either SUV membranes or SUPER templates was sufficient to block the interaction between SUVs and GUVs labeled with complementary oligos (SUVs with strand A and SUPER templates with strand A'). SUVs and SUPER templates both labeled with surface-bound $PEG_{5K}$ showed no membrane interaction. Scale bar: 10 μm.

### Supplementary Information

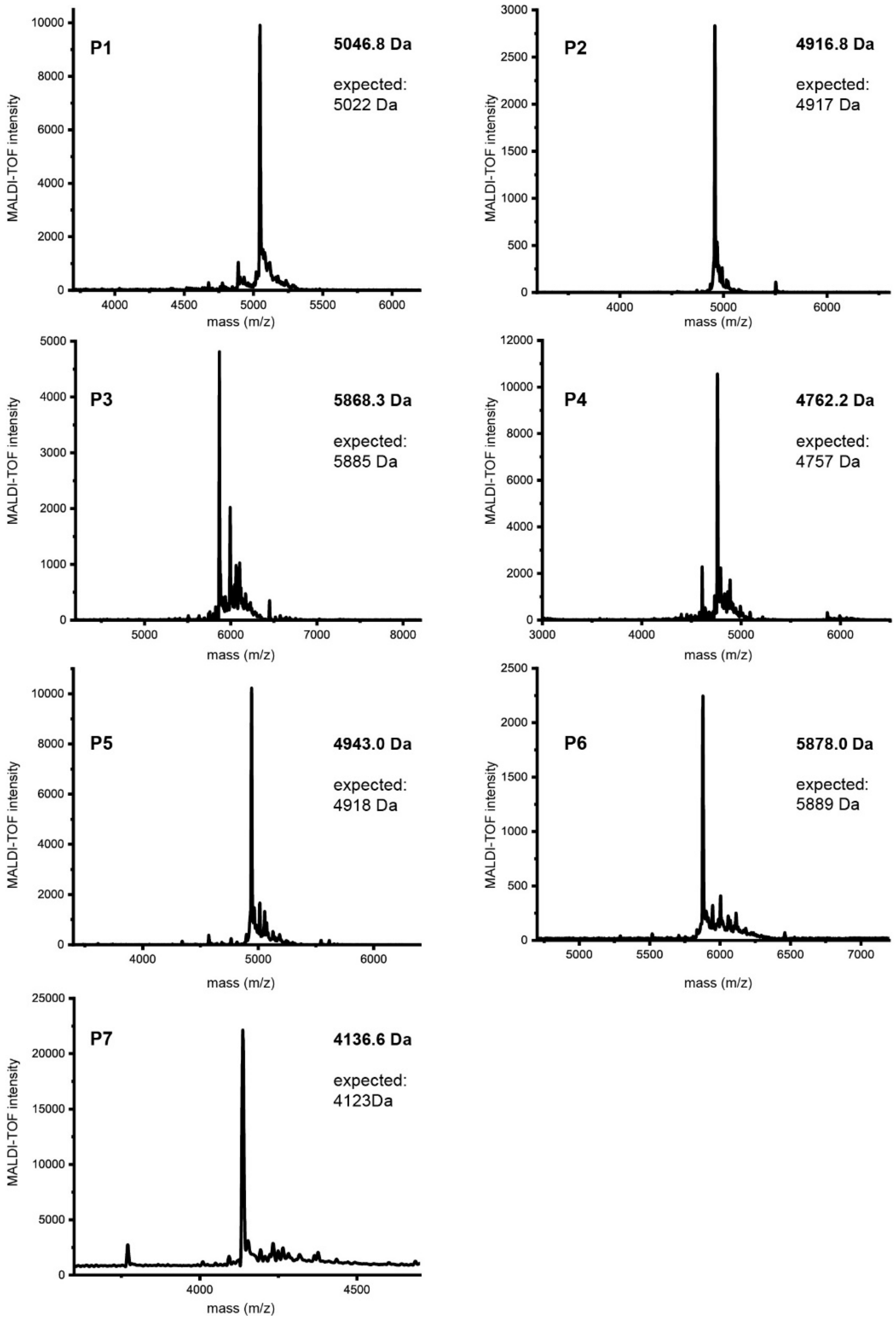


**Figure S3. MALDI-TOF MS spectra of indicated peptides.** Peaks of synthesized peptides mentioned in Table S3 match the expected molecular weights. See Table S3 for specific details on peptides P1-7.

**Supplementary Information**

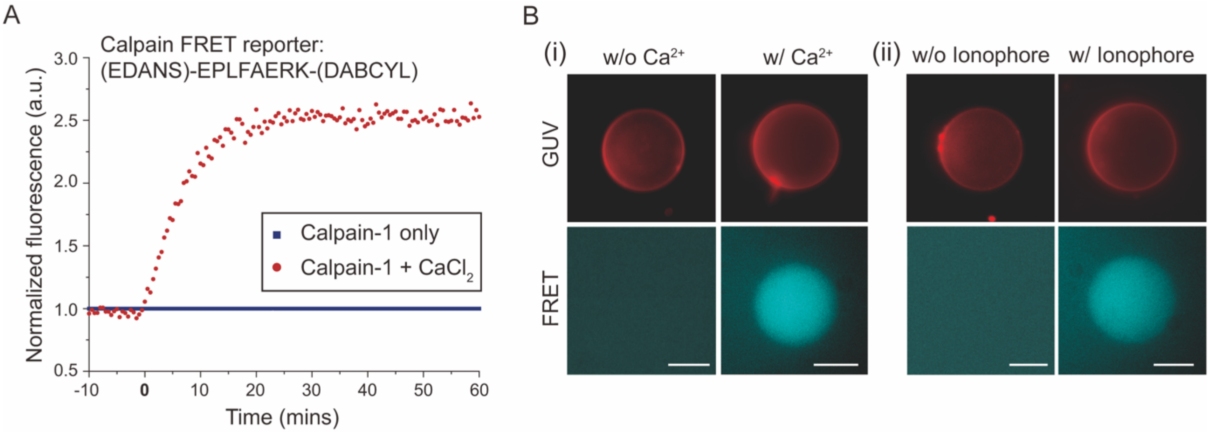


**Figure S4. Calcium-dependent calpain-1 activity tested by EDANS/DABCYL FRET pair**. (A) Time course of the calpain-1 activity of EDANS/DABCY FRET pair connected with calpain cleavage site by monitoring EDANS fluorescence by using a fluorescence plate reader. 5 mM $CaCl_2$ was added to the solution at T = 0 min. (B) Encapsulation of 250 nM calpain-cleavable EDANS/DABCYL FRET reporters inside GUVs. EDANS fluorescence was observed in GUVs co-encapsulating EDANS/DABCYL FRET pair and 3 mM $CaCl_2$ while GUVs without $Ca^{2+}$ inside exhibited low/no fluorescence (i). In the presence of 3 mM $CaCl_2$ in the outer solution, EDANS fluorescence was detected inside GUVs encapsulating EDANS/DABCYL FRET pair following the addition of 2 μM calcium ionophore A23187 in the outer solution (ii). Scale bars are 10 μm.

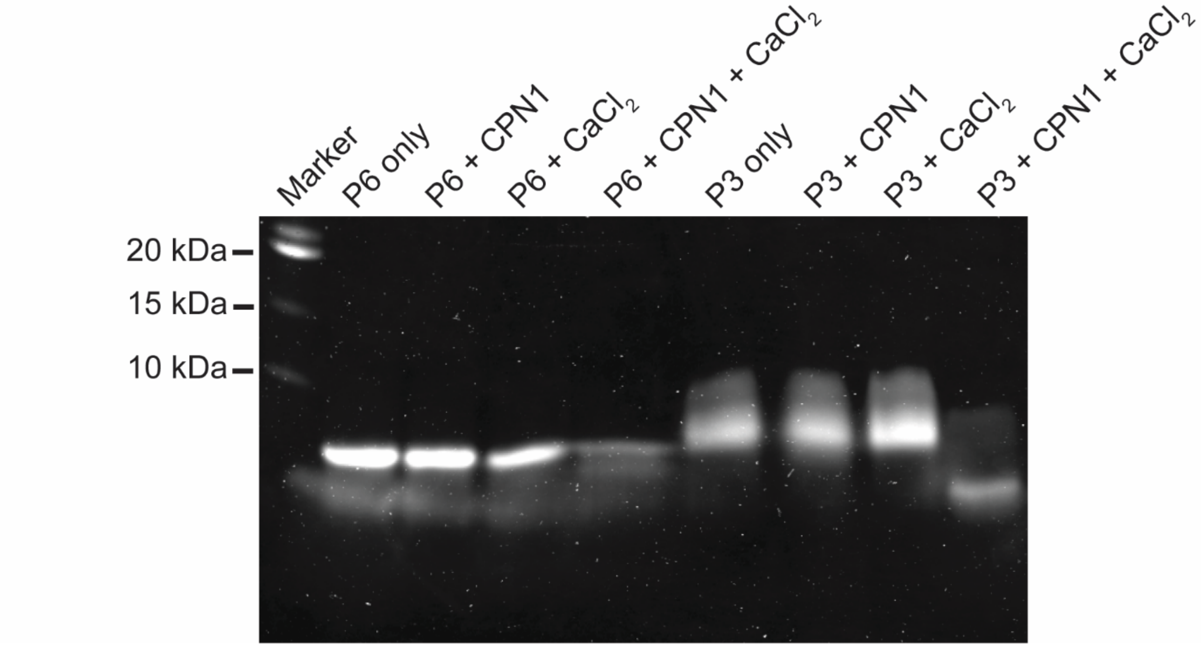


**Figure S5. Sodium dodecyl sulfate–polyacrylamide gel electrophoresis (SDS-PAGE) analysis to validate the calpain cleavage site (CCS) in bulk reaction.** For peptides P3 & P6, which contain CCS between peptide K or E and $PEG_{24}$ (around 1 kDa), only the addition of both 450 nM calpain-1 and 3 mM $CaCl_2$ in bulk reaction resulted in a significant decrease in the size of peptide, which suggests cleavage of the peptide at the CCS.

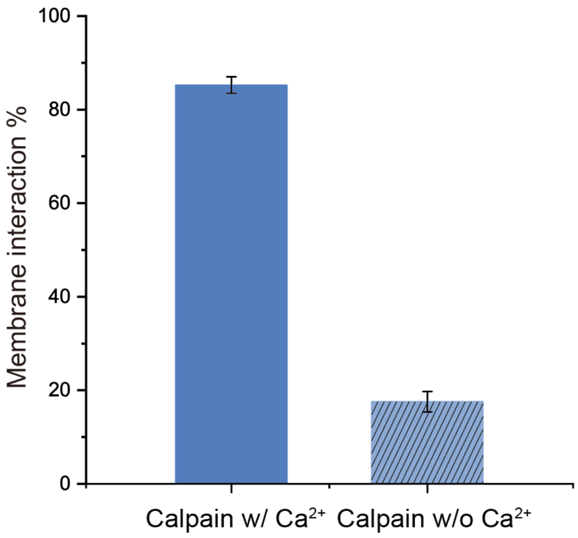


**Figure S6. Quantification of calcium-activated membrane interactions between SUPER templates and SUVs mediated by DNA oligonucleotides.** SUPER templates labeled with NBD-PE are functionalized with DNA strand A while SUVs labeled with Rhod-PE are functionalized with DNA strand A'. In the presence of 225 nM calpain-1, the bar graphs display the percentage of lipid-coated beads showing membrane interaction with SUVs by the red fluorescence detected on SUPER templates membranes with or without the addition of 5 mM $CaCl_2$. At least 60 lipid-coated beads were analyzed for each condition. All experiments were repeated three times under identical conditions. The error bars represent standard errors.

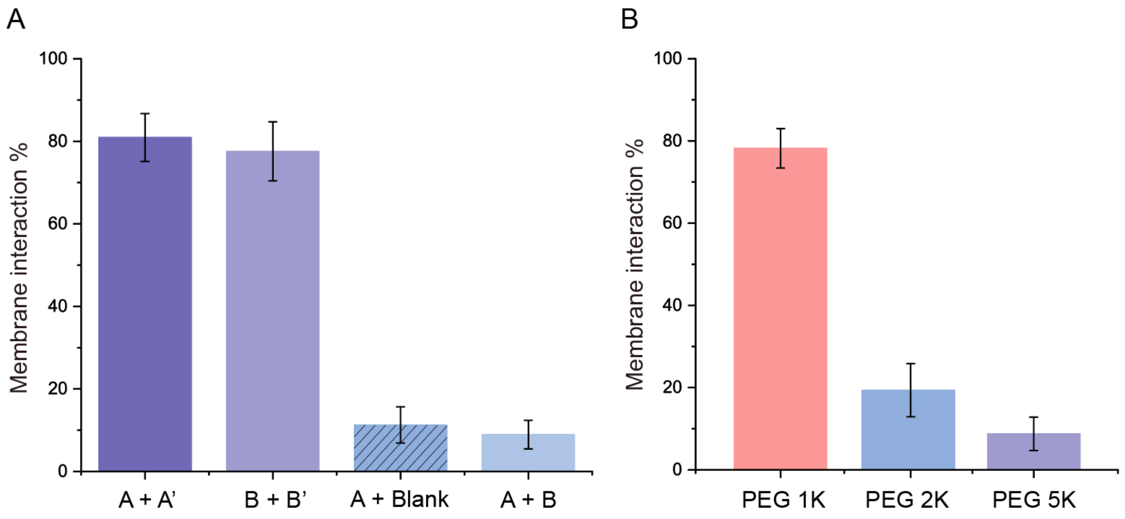


**Figure S7. Quantification of DNA-mediated membrane interactions between GUVs and SUVs.** (A) GUVs labeled with NBD-PE are functionalized with DNA strand A or B respectively while SUVs labeled with Rhod-PE are functionalized with DNA strand A', B', or no DNA. The bar graphs represent the percentage of GUVs showing membrane interaction with SUVs by the red fluorescence GUV membranes. (B) Both GUVs and SUVs membranes were covered with complementary DNA and PEG chains. The bar graphs display the percentage of GUVs showing red fluorescence rings to the total number of GUVs. At least 30 vesicles were analyzed for each condition. All experiments were repeated three times under identical conditions. The error bars represent standard errors.

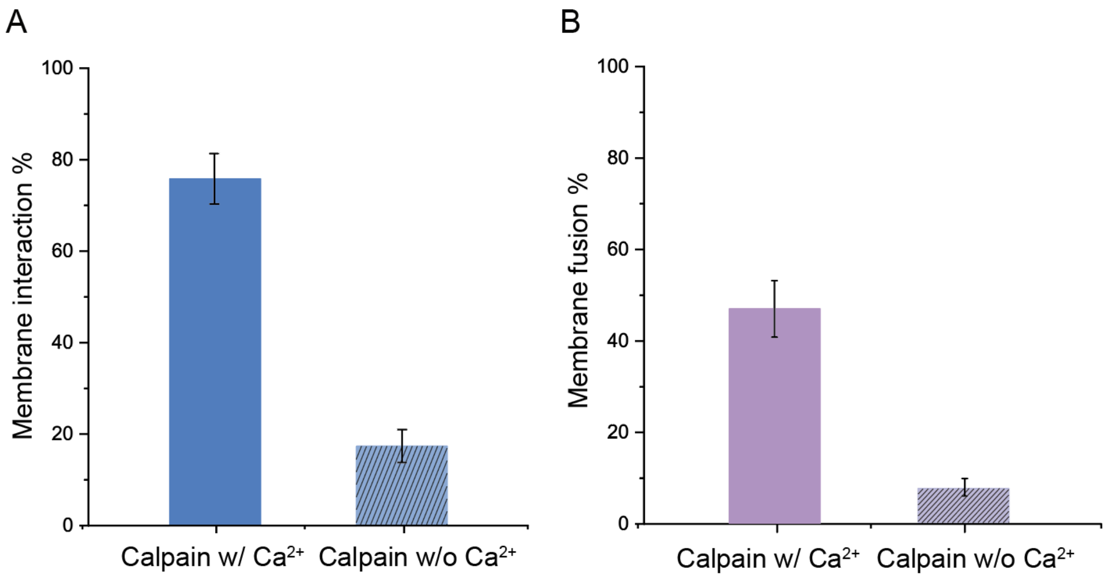


**Figure S8. Quantification of calcium-activated membrane interactions and membrane fusions between GUVs and SUVs mediated by DNA oligonucleotides.** GUVs labeled with NBD-PE were covered with DNA strand A and calpain-cleavable $PEG_{2k}$ while SUVs labeled with Rhod-PE were covered with DNA strand A' and calpain-cleavable $PEG_{2k}$. (A) The blue bars represent the percentage of GUVs showing membrane interaction with SUVs by the red fluorescence on GUV membranes. (B) The purple bars represent the percentage GUVs showing membrane interactions with SUVs that also show GUV membrane fusion with SUVs. At least 30 vesicles were analyzed for each condition. All experiments were repeated three times under identical conditions. The error bars represent standard errors.

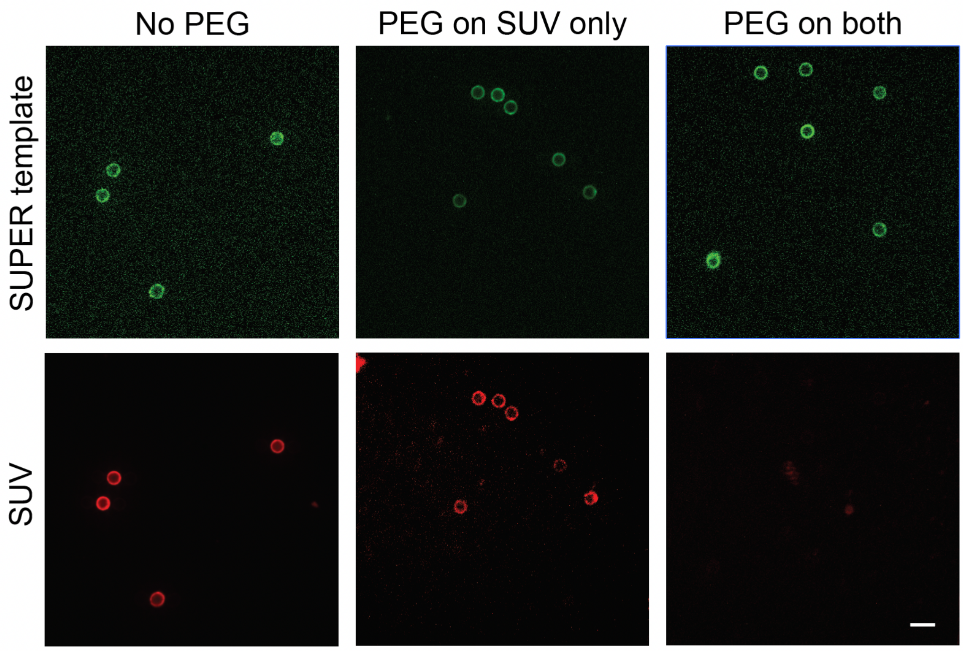


**Figure S9. Inhibition of peptide-mediated membrane interactions with surface-bound PEG$_{2K}$.** SUPER templates and SUVs decorated with membrane-bound peptides K (peptide P1) and E (peptide P1), respectively. Membrane interactions were observed when no surface-bound PEG$_{2K}$ polymers were present (No PEG) and when PEG$_{2K}$ polymers were only on the SUV membrane (PEG on SUV only). When PEG$_{2K}$ chains are conjugated to both membrane surfaces, no membrane interactions are seen. Scale bar: 10 µm.

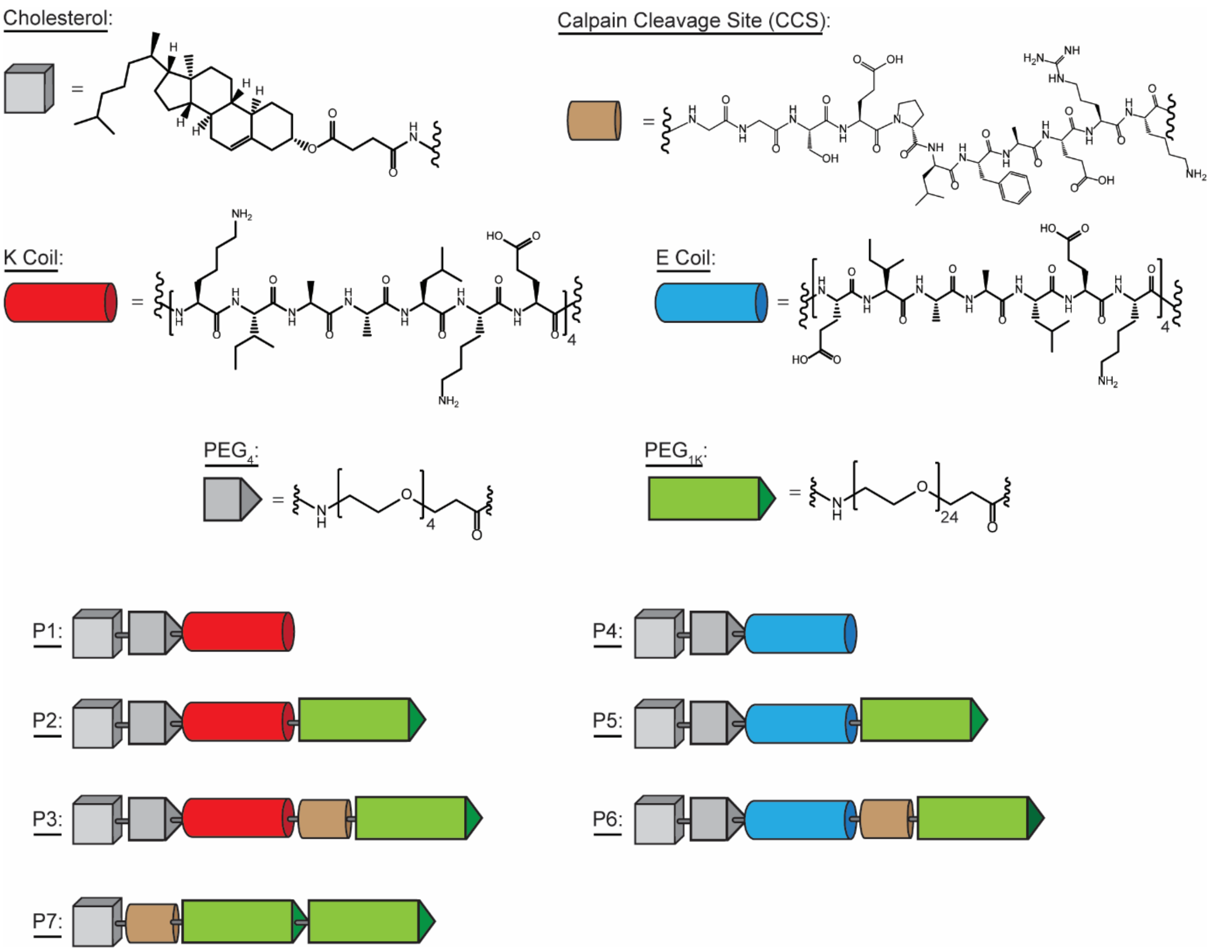


**Figure S10. Schematics of synthesized peptides.** All seven peptides synthesized are graphically represented above. This figure complements the peptide details listed in Table S1.

### Supplementary Information

### Tables

**Table S1. Synthesized peptides**

| Peptide | Description | Sequence | Molecular Weight (Da) |
|---|---|---|---|
| P1 | Cholesterol-$PEG_4$-K Coil | Cholesterol-$(PEG)_4$KLNKWWYKRKELAAIEKELAAIEKELAAIEKELAAIK | 5022 |
| P2 | Cholesterol-$PEG_4$-K Coil-$PEG_{24}$ | Cholesterol-$(PEG)_4$KIAALKEKIAALKEKIAALKEKIAALKE$(PEG)_{24}$ | 4917 |
| P3 | Cholesterol-$PEG_4$-K Coil-CCS-$PEG_{24}$ | Cholesterol-$(PEG)_4$KIAALKEKIAALKEKIAALKEKIAALKEEPLFAERK$(PEG)_{24}$ | 5885 |
| P4 | Cholesterol-$PEG_4$-E Coil | Cholesterol-$(PEG)_4$KKRRAKSQEKLAAIKEKLAAIKEKLAAIWEKLAAIK | 4757 |
| P5 | Cholesterol-$PEG_4$-E Coil-$PEG_{24}$ | Cholesterol-$(PEG)_4$EIAALEKEIAALEKEIAALEKEIAALEK$(PEG)_{24}$ | 4918 |
| P6 | Cholesterol-$PEG_4$-E Coil-CCS-$PEG_{24}$ | Cholesterol-$(PEG)_4$EIAALEKEIAALEKEIAALEKEIAALEKEPLFAERK$(PEG)_{24}$ | 5925 |
| P7 | Cholesterol-CCS-$PEG_{48}$ | Cholesterol-GGSEPLFAERK$(PEG)_{48}$ | 4123 |

* Block diagram of peptides can be found in Figure S10.

**Table S2. GUV lipid compositions**

| Experiment | Lipid composition for GUVs (molar ratio) |
|---|---|
| Membrane interaction w/o Chol-CCS-PEG | DOPC:DOPE:cholesterol:NBD-PE = 74.9: 5: 20: 0.1 |
| Membrane interaction w/ Chol-CCS-PEG | DOPC:DOPE:cholesterol:NBD-PE:Chol-CCS-PEG2K = 69.9:5:20:0.1:5 |
| Membrane fusion w/o Chol-CCS-PEG | POPC:DOPE:NBD-PE = 94.9:5:20:0.1 |
| Membrane fusion w/ Chol-CCS-PEG | POPC:DOPE:NBD-PE:Chol-CCS-PEG2K = 89.9:5:20:0.1:5 |

**Table S3. SUV lipid compositions**

| Experiment | Lipid composition for SUVs (molar ratio) |
| --- | --- |
| Membrane interaction w/o Chol-CCS-PEG | DOPC:DOPE:cholesterol:Rhod-PE = 69.9:10:20:0.1 |
| Membrane interaction w/ Chol-CCS-PEG | DOPC:DOPE:cholesterol:Rhod-PE:Chol-CCS-PEG2K = 64.9:10:20:0.1:5 |
| Membrane fusion w/o Chol-CCS-PEG | DOPC:cholesterol:Rhod-PE:DGS-NTA (Ni) = 34.9:15:0.1:50 |
| Membrane fusion w/ Chol-CCS-PEG | DOPC:cholesterol:Rhod-PE:DGS-NTA (Ni):Chol-CCS-PEG2K = 34.9:10:0.1:50:5 |
